# PCR–based assays for determining mating status in field–weathered *Ceratitis capitata* with enhanced precision across conventional, quantitative, and droplet digital platforms

**DOI:** 10.64898/2026.08.12.744474

**Authors:** JAP Marcelino, CJ Zuck, H Urbina, MR Moore, MS Siderhurst, A Hurst, K Fairbanks, J Stanley

## Abstract

Accurately determining the mating status of the agricultural fruit fly pest *Ceratitis capitata,* commonly known as Medfly, is essential for timely and effective eradication efforts. To overcome the limitations of subjective DAPI–based staining assessments of females captured in Jackson dry traps and Multilure liquid traps, we developed a multi–tier molecular diagnostic method that unequivocally detects mating status using DNA probes targeting the male–specific Y114 locus on the Y-chromosome of the species. Our protocol integrates morphological evaluation with increasingly sensitive molecular assays through the following steps: 1) A preliminary quality assessment of the specimens’ physical condition, DNA preservation, and mating status using conventional PCR followed by agarose electrophoresis (cPCR); 2) Quantification and real–time detection of sperm presence via quantitative PCR (qPCR); and 3) Detection of trace sperm amounts through droplet digital PCR (ddPCR). This PCR–based framework is designed for samples collected in the field, enabling accurate analysis of specimens exposed to adverse environmental conditions and varying levels of preservation after 2- and 3-weeks weathering times in traps. It allows quantitative determination of mating status even when sperm concentrations are extremely low, such as during transient copulation, and achieves detection limits down to approximately 14 spermatozoa in a mated female. By accounting for variable specimen quality and the performance characteristics of each molecular platform, this tiered approach ensures highly sensitive and unequivocal detection of mated females. The methodology can be used to assist eradication efforts across the *C. capitata* geographic range through the timely detection of mated females, halting their expansion and establishment into novel regions reducing control and eradication costs.

## 1 Introduction

Exotic true fruit flies (Diptera: Tephritidae) pose a threat to the agricultural industry worldwide, threatening food security and the economy. The fruit and vegetable industry in the United States, valued at $48 Billion (Huang et al., 2022), is particularly vulnerable to these pests. Globally, one of the most destructive species in the agricultural industry is the Mediterranean fruit fly (Medfly), *Ceratitis capitata* (Wiedemann), with an estimated 408 suitable host plants and fruits (Liquido et al., 2020). Recurrent Medfly detections and subsequent eradications have occurred in Florida from 1929 to 1998, and more recently from 2010 to 2011 (Denmark, 1956; Halbert, 2010; Thomas et al., 2022). Prevention and eradication programs rely on the capture performance of traps and lures (IAEA, 2003) established across detection grids such as the 56,000 traps spanning over 20,720 km^2^ in Florida’s Fruit Fly Exclusion and Detection Program (Kalaris et al., 2014; Steck et al., 2019). A critical aspect of prevention and eradication programs is the accurate detection of the mating status of females since the presence of such specimens indicates the potential of a breeding population and subsequent expansion. In the United States, the detection of a single mated female triggers an emergency delimitation, eradication, and quarantine protocol (USDA-APHIS, 2020), increasing trap density and monitoring schemes from the detection point to a 10 miles radius, with 2050 traps in an 81 sq mile area for *C. capitata* (FDACS-USDA, 2013). Emergency response protocols become very costly when multiple mated Medfly females are detected. For example, eradication efforts in 1997 reached a total cost of $25 million (USDA, 2015; Szyniszewska et al., 2016).

Currently DAPI-staining (4′,6-diamidino-2-phenylindole) based procedures are used to detect sperm in the female fertilization chamber the ventral receptacle, as well as sperm storing spermathecae (Marchini et al., 2001; Shadmany et al., 2021; Pérez-Staples et al., 2023). DAPI DNA staining techniques are time-consuming, inconsistent and subjective as they rely on the visual assessment of stained spermatozoa. DAPI staining of spermatozoon nuclear protamines can lead to incorrect assessments due to extremely compact DNA in the nucleus of the spermatozoa and other nuclear structures with heterochromatic DNA, especially in the Y-chromosome (hereafter as Y-chrom) of *C. capitata* (Zhou et al., 2000), which hinders the penetration capability of the stain, thus creating uneven staining (Fuller, 1993). The clusters of elongated needle-shape spermatozoa in *C. capitata* (see image in Twig and Yuval, 2005) can appear structurally similar to other anatomical features in the fly. In addition, the mating behavior of intermittent copula (Prokopy and Hendrichs, 1979) may translate into very few spermatozoa being transferred at a single given mating event, making the visual observation of staining difficult. Moreover, this problem is further exacerbated by the fact that monitoring schemes for traps in the field can be scheduled at a maximum of three weeks intervals (FDACS-USDA, 2013), promoting specimen deterioration in open-air as in the Jackson dry traps, or liquid traps as in Multilure McPhail traps, which can severely compromise spermatozoa and their staining. At 3.5-5 mm, deterioration is particularly important for *C. capitata* which body is at the lower size range for Tephritidae flies (Weems, 1981). Small flies are very prone to decay given their small internal mass (Kühsel et al 2017). In addition, DAPI is a known mutagen and its use requires protective equipment to avoid contact with skin (Stauffert et al., 1990). The need for an unambiguous and quantitative approach which can detect spermatozoa in inseminated females persist.

*Ceratitis capitata* has an XX-XY sex determination system with heterogametic XY males and homogametic XX females (Radu et al., 1975). The male-determining region of the Y-chrom of *C. capitata* was previously identified by Willhoeft and Franz, 1996. Presence of the Y-chrom in a female will confirm fertilization. Male *C. capitata* Y-specific amplicons have been previously identified (Anleitner and Haymer, 1992; Zhou et al. 2000; Douglas and Haymer, 2004) but not tested for screening the mating status of females. The authors detected and used the male-specific amplicon of the Y-chrom for molecular sexing under optimal conditions with ethanol preserved specimens. Another approach attempted to estimate stored sperm in female *C. capitata* (San Andrés et al 2007, Catalá-Oltra et al., 2020), but these studies were also designed using optimal conditions, utilizing specimens preserved in ethanol, directly placed in the preservative liquid, or from 1-week weathered field traps. These experiments provided results on number of sperm in spermatheca of *C. capitata* based on conventional PCR (cPCR) and quantitative PCR (qPCR) but did not consider the progressive decay of the specimens in field traps which as mentioned can remain in field conditions up to three weeks (FDACS-USDA, 2013; FAO/IAEA, 2018). Similar proof-of-concept approaches in laboratory conditions were developed for the Queensland fruit fly, *Bactrocera tryoni* (Froggatt) with the same limitations. Moreover, for *C. capitata*, the qPCR technique uses a SYBR^®^ dye-based detection which presents the following limitations: Low specific binding to the target amplicon, thus increasing the chance of false positives; no multiplexing capability which limits the use of controls in duplex qPCR reactions, especially in the context of decayed specimens where an internal control for genetic integrity of the specimen in the reaction is needed; and inhibition of PCR at high concentrations (Gudnason et al., 2007).

We aim to develop a three-tier genetic approach with increasing detection and sensitivity to low-abundance Y-chrom present in mated female *C. capitata* using standard end-point conventional PCR (cPCR) and gel electrophoresis (Tier 1), real-time PrimeTime® double-quenched high target-specific probes for quantitative PCR (qPCR) (Tier 2), and droplet digital PCR (ddPCR) (Tier 3). Analytical and sample performance of synthetic DNA target amplicons and, more importantly, field specimens from all possible scenarios (i.e., collected at 2- and 3-week intervals from Jackson and McPhail-like traps with highly variable sperm presence) were evaluated. This tier approach presents the following advantages: **1.** Laboratories with different technical capabilities can detect mating status of female *C. capitata* using the presence/absence of the Y-chrom, from a cost-and-ease gel-based electrophoresis protocol to robust real-time PCR and ddPCR protocols; **2.** Enhanced resolution power, sensitivity and specificity for detecting target amplicons of the Y-chrom at increasingly lower concentrations from Tier 1 to Tier 3; **3.** The capability to process specimens at different stages of decay since specimens will be exposed to open air conditions or H_2_O and propylene glycol based preservation agents in traps, collected at a maximum of 3-weeks from capture; **4.** PrimeTime^®^ double-quenched target-specific probe-based qPCR will ensure assay sensitivity and detection capability of the target amplicon, as well as overriding the limitations described for SYBR^®^ dye-based assays; **5.** Mating status confirmation can be done in a few hours, thus allowing for prompt implementation of control measures based on robust genetic detection of Y-chrom amplicons; and **6.** The techniques can be easily implemented to facilitate the analysis of field-collected trap specimens in contrast to more laborious full genome genotype-by-sequencing approaches or current DAPI-staining based procedures.

## 2 Materials and Methods

### 2.1 *C. capitata* sample collection and weathering times

A total of 105 specimens were assayed, which included cohorts of males (♂), unmated or virgin females (*v*♀), and mated females (*m*♀) originating from colony populations in distinct geographic locations (Supplementary Table 1A-1C) of Argentina (N=45, ♂=15, *m*♀=15, *v*♀=15) and Hawaii, United States of America (N=45, ♂=15, *m*♀=15, *v*♀=15). In addition, we also obtained sterile irradiated males (*s*♂, N=15) of the Genetic Sexing Strain (GSS) Vienna 8^D53-^ from the El Pino mass-rearing facility in Guatemala which utilizes Sterile Insect Technique (SIT) to produce sterile male *C. capitata* for eradication and control programs of Guatemala, Mexico, and the United States of America (Ramírez-Santos et al., 2025).

In Argentina and Hawaii where populations of Medfly are considered established, mating status of females was determined by sexing the pupae and capturing females upon emergence. Virgin females were kept separated, while *m*♀ were placed in cages with mating arenas and were observed to engage in non-interrupted copula for at least 40 minutes (min). This observational period is essential given the intermittent copulation behavior of the species (Prokopy and Hendrichs, 1979). Specimens were placed directly in 95% ethanol (hereafter EtOH) (quality control specimens) or freeze-dried and stored at -20°C prior to shipment (field weathered specimens). Samples were then shipped to the Florida Department of Agriculture and Consumer Services, Division of Plant Industry (FDACS-DPI) in Gainesville, Florida, the United States of America.

At FDACS-DPI the specimens in EtOH were used as quality control for direct DNA extraction, while the freeze-dried specimens were placed in two types of traps:

1. Jackson® traps, hereafter JK, (Harris et al., 1991), a dry-based trap in an open-air field setting with sticky inserts to replicate the physical decay of specimens captured in these traps which are used in area-wide detection grids worldwide (Shelly et al., 2014; Gilbert et al., 2013, Kalaris et al., 2014; Steck et al., 2019; Perez-Staples et al., 2020). Specimens were transferred with flame-sterilized stainless-steel forceps onto individual JK traps in three replicates per specimen type as ♂, *s*♂, *v*♀, and *m*♀, and per geographic origin.
2. Multilure® traps, hereafter ML, a liquid-based McPhail-like trap (FAO/IAEA, 2018), also a key component of monitoring grids for prevention, surveillance, and eradication of fruit flies. ML traps utilize a liquid preservative, typically a 10 % propylene glycol (PG)-based solution where flies are collected, with an attraction lure to induce fruit fly entry into the trap (Martinez et al., 2007; FDACS-USDA, 2013; FAO/IAEA, 2018; Perez-Staples et al., 2020). Propylene glycol (PG) was sourced from commercially available automotive antifreeze/coolants currently used across the state-wide fruit fly monitoring grid of Florida (Arctiq® Eco Safe Green, or DefendAL® Enviro). Antifreezes were analyzed for their PG content with High Resolution-Accurate Mass (HRAM) GC-MS chromatography at the Mass Spectrometry Research and Education Center of the University of Florida; their content was determined to be 98% PG. Specimens were transferred with flame-sterilized stainless-steel forceps into containers with 400 mL of 10% Arctiq solution (90:10 H_2_ O:Arctiq) and allocated into a Percival^®^ I-30VL incubator (Percival Scientific, Perry, IA) with controlled temperature (28 ± 2 °C), relative humidity set at 70 % and photoperiod of LD 12:12 h to mimic field environmental conditions.

Specimens remained in JK and ML traps for two weeks, which is the standard time fruit flies remain in traps in current field monitoring schemes of Florida. Additional specimens remained for the maximum trap retrieval time of three weeks. Please refer to Supplementary Table 1 for specimen characteristics and weathering times for JK and ML traps.

### 2.2. *C. capitata* genomic DNA (gDNA) extraction

Individual specimens were retrieved with sterile forceps from their traps and were processed accordingly prior to abdomen excision as:

1. Control specimens in EtOH, removed from their collection tube and were air-dried on a KimWipe for 5 min.
2. JK trap specimens, individually submerged in 1.5 mL microcentrifuge tubes filled with 1 mL of Histo-clear II (a non-toxic solvent that does not impact DNA yield) for 15 min, then rinsed three times with EtOH, and placed on a KimWipe to air-dry for 5 min.
3. ML trap specimens, placed in a sterile Petri dish of EtOH to soak for 5 min, then placed on a KimWipe to air-dry for 5 min.

For traps with heavy debris or non-target insect presence, we recommend specimens be surface sterilized after collection utilizing a 45 seconds (s) submersion in a 0.5% sodium hypochlorite solution, followed by a triple rinse with DNase and RNase free ultra-pure water (hereafter hpH_2_O). This helps reduce contaminants that may interfere with PCR efficiency or gel visualization.

Once air-dried, specimens were moved to a sterile Petri dish and their abdomens were excised with a one-time use disposable scalpel (Exel International Inc., item #14-840-00). Remaining tissues were collected in EtOH and stored at the FDACS-DPI Methods Arthropoda collection.

Abdomens were individually placed in sterile 2 mL screw-cap tubes containing one 3.2 mm ⌀ stainless steel bead (Benchmark Scientific Inc., Sayreville, United States, item #D1134-50) and stored at -20 °C overnight. Frozen abdomens were then pulverized using a D2400Homogenizer (Benchmark Scientific Inc., Sayreville, United States) for 15 s at a speed of 5 M/s to achieve uniform disruption of specimen tissues. Tubes were then vortexed at 3000 rpm for 10 s and briefly centrifuged. After, 180 µL of buffer ATL (Qiagen GmbH, Germany) and 20 µL of Proteinase K were added to the samples, then briefly vortexed and centrifuged. Samples were placed in a water bath at 56 °C for overnight incubation to better cleave peptide bonds and break down proteins. After overnight incubation, an additional 10 µL of Proteinase K was added to the tubes, and the temperature was increased to 62 °C for 4 h. Subsequent DNA extraction steps were performed using the DNeasy^®^ Blood & Tissue Kit (QIAGEN GmbH, Hilde, Germany) per the manufacturer’s protocol. Final elution of DNA in 31 µL hpH_2_O.

Total gDNA was quantified using a Qubit™ dsDNA 1x HS Assay Kit with an Invitrogen™ Qubit™ 4 Fluorometer following the manufacturer’s protocol. Total gDNA was stored at -20 °C prior to downstream processing.

### 2.3. Primer design for Tier 1 (cPCR), Tier 2 (qPCR), and Tier 3 (ddPCR)

A unique male specific and repetitive A-T nucleotide rich Y-chrom amplicon, Y114 (Genbank accession no. AF071418) was previously identified by Anleitner and Haymer (1992), Zhou et al. (2000) and Douglas and Haymer (2004). We contrasted the Y114 sequence with possible matches in NCBI GenBank nucleic acid sequence databases using the National Center for Biotechnology Information Basic Local Alignment Search Tool, BLAST (Altschul et al 1990). Observed matches were limited to *C. capitata*, and included those generated by the above authors in addition to Y114 (GenBank accession nos. AF071418, AF115330, AF115331, AF154063). In addition, GenBank accession no. GU122240 from a gene expression embryogenetic study for sex in *C. capitata* embryos (Gabrieli et al., 2010), and a computational predicted enzyme in *C. capitata* were also listed (GenBank accession no. XM023303525).

The 1405 bp Y114 amplicon poses challenges for DNA amplification given the unusually high A-T content of AT’s repetitive elements reaching 83% (Zhou et al. 2000). This is further complicated given the AT’s-homopolymer of ∼200 bp is the male specific fragment within the male specific amplicon. Taking these parameters into consideration and given qPCR and ddPCR technology preferentially sequence short amplicons (≤150-200 bp) (Rocha et al., 2016; Nolan et al 2013; Martin et al 2013; Abellan-Schneyder et al., 2021; Van Holm et al., 2021; Lim et al., 2021) we used the forward primer designed by Zhou et al. 2000 for the Y114 AT-element (Y114F10, 5’– TGCCAAAGCACTATCTTCGGAAG –3’) and designed a new reverse primer (Y-RV2, 5’– GGCTAATTGTTTTACCATAAC –3’), that includes the 200 bp AT’s-homopolymer within the AT-element. We did not use but Zhou et al. 2000 reverse primer (Y114B13), given this combination generates a 710 bp amplicon not suitable for qPCR and ddPCR. Given the highly rich AT composition of the Y114 amplicon and the need to include GC nucleotides in primer design to increase binding stability given GC have three hydrogen bonds (G☰C) whereas AT have two (A═T), our reverse primer had to be the closest adjacent sequence to the AT element with these characteristics.

The combination of these primers generated a fragment of 253 bp of Y-chrom with A-T nucleotides accounting for 73% of its content. Smaller amplicons would include even higher A-T content which would compromise PCR efficiency and were therefore not viable.

In addition, we include in all individual reactions, across all tiers, a control gene to attest for the physical and genetic integrity of the specimens, following Zhou et al. 2000. We used the muscle-specific *Actin* gene CcA1 (hereafter *Actin*), a commonly used housekeeping gene in PCR studies with stable expression across development stages, which expresses as a protein involved in multiple functions of invertebrates such as muscle contraction, cell division and differentiation (He & Haymer, 1993; Hightower & Meagher, 1986; Ponton et al., 2011; Rahman et al., 2021). Therefore, all tiers in our methodology are comprised of duplex reactions with Y-chrom and *Actin* gene amplicons.

Following the same premises listed above for cPCR and qPCR, as well as primer design, the amplicon selected comprises a modified Zhou et al. 2000 forward primer, as *Actin*-fw-Set4 5’– CTTGGACTT TGAGCAGGAGATG –3’, and a new reverse primer, as *Actin*-rv-Set4 5’– AGGAATGAGGGCTGGAAGA –3’, generating an amplicon of 141 bp with a 39 % A-T content.

The efficacy of this combination of primers and amplicon sizes in duplex reactions warranted high reaction efficacy across the three tier protocols for mating status detection in female *C. capitata*; however, for Tier 1 (cPCR) we locked nucleic acids in the reverse Y-chrom primer (+, denoted in bold and in front of the locked nucleic acid) to increase its specificity to the short target amplicon and avoid non-specific bands in gel electrophoresis visualization (as Y-RV2-lkd 5’ – GGCT**+**AATT+GTT+TTA+CCA+TAAC -3).

### 2.4. Probe design for qPCR and ddPCR

To enhance amplification fidelity and precision to our target Y-chrom and *Actin* amplicons, we used PrimeTime^®^ double-quenched high-fidelity probes for qPCR and ddPCR. Double-quenched probes include an additional internal quencher in the probe decreasing the distance between the 5′ fluorophore and quencher. This additional quencher translates into more robust quenching and lower background noise, as well as earlier detection of target amplicons.

We used PrimeTime® Double-Quenched Probes from Integrated DNA Technologies Inc. (IDT) in 100 µM concentrations, which contain a 5′ fluorophore (FAM reporter dye for the Y-chrom amplicon and SUN reporter dye for the *Actin* amplicon), a 3′ Iowa Black^®^ quencher, and an internal ZEN™ quencher.

### 2.5 Protocol for Tier 1 (cPCR) assay

Duplex PCRs (25 μL final volume) consisted of one Lumiprobe DryDrops® HotStart Lyophilized PCR Bead (Lumiprobe Corp., item # 21534), 20 μL hpH_2_O, 0.5 μL of 5 µM *Actin* Fw, 0.5 μL of 5 µM *Actin* Rv, 1 μL of 10 µM Y-chrom Fw, 1 μL of 10 µM Y-chrom Rv, and 2 μL of genomic DNA template. *Actin* is over-expressed across all tissues (Rahman et al., 2021) hence we adjusted the molarity to balance the duplex reaction with the Y-chromosome to avoid preferential sequencing of *Actin* in reaction.

Reactions were run on an Applied Biosystems MiniAmp™ Plus Thermal Cycler (ThermoFisher Scientific, Bedford, United States) with the following cycling conditions: 95 °C for 2 min followed by 34 cycles of 95 °C 30 s; 60 °C 25 s, with no final extension step, and a ramp rate of 2.5 °C/s. No template reactions (NTCs) were used as negative controls.

Gel electrophoresis utilized 2 % agarose in Tris-borate-EDTA (TBE) gel with SYBR Safe Gel Stain run at 110 V for 60 min. Wells were loaded with 20 μL of PCR product and 4 μL of 6× TriTrack DNA Loading Dye alongside 2 μL of GeneRuler 100 bp DNA Ladder. Gel bands were visualized using a Bio-Rad GelDoc Go Imaging System with UV/Stain-Free Tray (Table 1 for summarized description of protocols).

**Table 1.**
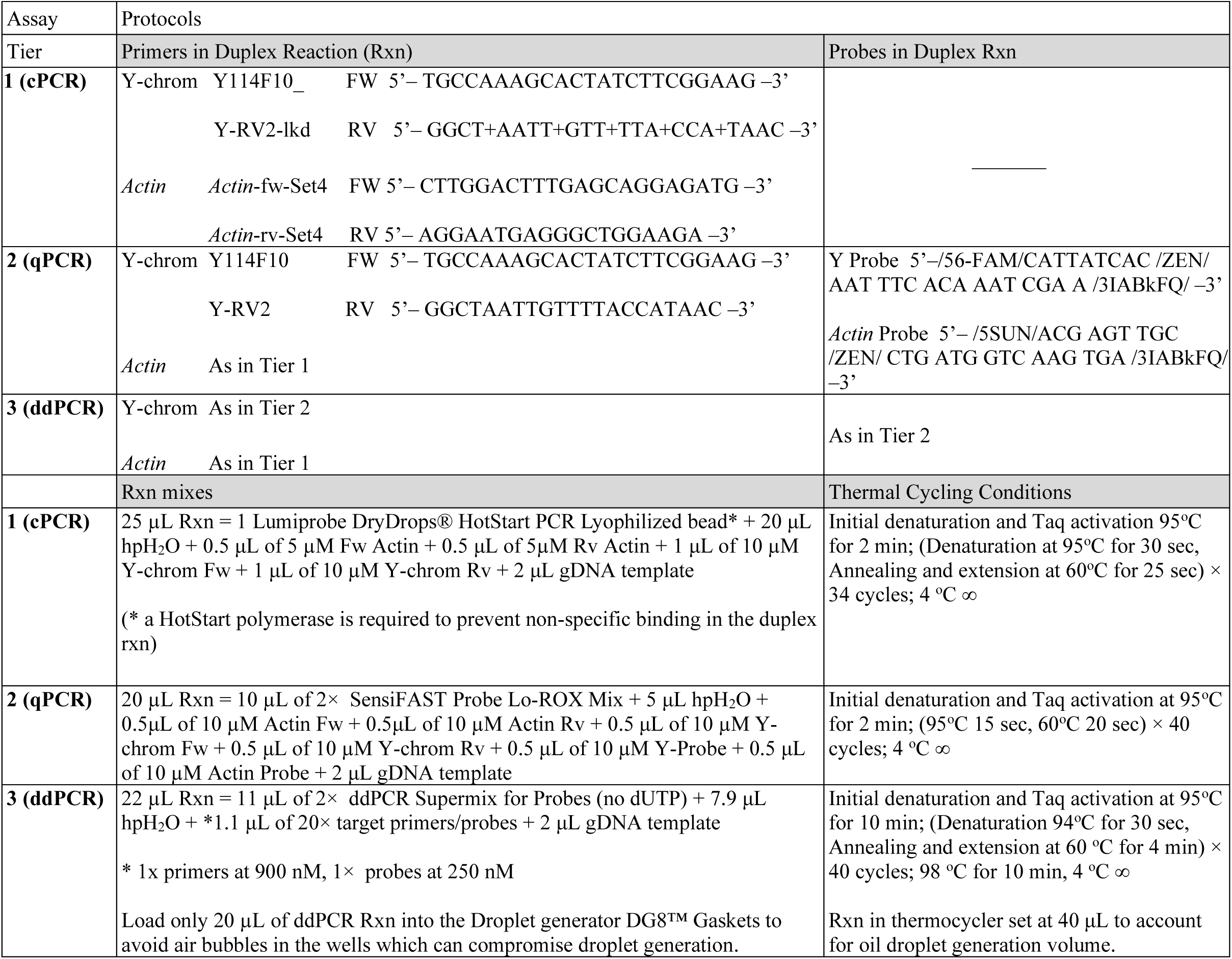
Protocols for assay methodology per reaction (Rxn). FW=primer forward, Rv=primer reverse.

### 2.6. Protocol for Tier 2 (qPCR) assay

Duplex qPCR reactions (20 μL final volume) consisted of 10 µL SensiFAST Probe Lo-ROX Mix (2×), 5 µL of hpH_2_O, 10µM of primers and probes at 0.5 µL volume, i.e., 0.5 µL *Actin* Fw, 0.5 µL *Actin* Rv, 0.5 µL Y-chrom Fw, 0.5 µL Y-chrom Rv, 0.5 µL Y-chrom Probe, 0.5 µL *Actin* Probe, and 2 μL of genomic DNA template. All reagents and reactions were maintained on Fisherbrand^TM^ Cooling Blocks during preparation.

Samples were run on an Applied Biosystems QuantStudio™ 5 Real-Time PCR System (ThermoFisher Scientific, Waltham, United States) with the following cycling protocol: 95 °C for 2 min followed by 40 cycles of 95 °C 15 s; 60 °C 20 s, with no final extension step, and a ramp rate of 2.5 °C/s. A variable number of no template control reactions (NTCs) were used as negative controls per assay sessions. Results were visualized using Thermo Fisher™ QuantStudio™ Design and Analysis 2 (DA2) software (Table 1 for summarized description of protocols).

Quality Control for qPCR followed the MIQE guidelines (Minimum Information for Publication of Quantitative Real-Time PCR Experiments) for replication and reproducibility (Taylor et al., 2010; Bustin et al. 2009, 2025a). Standard curves were generated using 10-fold serial dilutions (10^5^-10^0^ amplicon copies) of synthetic target *Actin* and Y-chrom sequences. We determined that concentrations above 10^5^ exceeded the upper limit of the dynamic range of the qPCR assay, which is also the recommended dynamic range of the QX200 ddPCR system we used (Bio-rad, 2014). Synthetic amplicon DNA targets were obtained from Eurofins Genomics Blue Heron LLC., Bothell, United States. The concentrated number of synthetic copies prior to 10-fold dilutions was 1.107×10^11^ per μL for the Y-chrom amplicon, and 4.685×10^12^ per μL for the *Actin* amplicon, per the manufacturer.

Synthetics were run in monoplex reactions for each amplicon, as well as duplex reactions to account for possible interaction between primers and probes in qPCR performance. Standard curves were derived from the serial dilution and the Quantification cycle (Cq) detection for each monoplex and duplex reactions. qPCR amplification efficiency and linear fitness were determined using a slope to efficiency calculator as Efficiency = -1+10^(−1/slope)^, and linear regression as Y=a-bX, where X is the slope of the regression line, and Y is the detection CT value. A slope of -3.322 indicates 100% efficiency, with target detection decreasing at ∼3.3 cycles between consecutive dilutions (Ma et al., 2020). Efficiency was calculated using the ThermoFisher Scientific qPCR Efficiency Calculator website interface, and regression fitness using JMP Pro v.19 software, Statistical Discovery LLC, Cary, NC, USA.

Sensitivity limits of Limit of Black (LoB), Limit of Detection (LoD) and Limit of Quantification (LoQ) were determined using three replicates per 10-fold serial dilutions from 10^-5^ to 10^0^ copies/μL for Y-chrom target amplicon and *Actin* housekeeping control gene, respectively.

Standard curves were used to evaluate these performance metrics. The Clinical Laboratory Standards Institute, CLSI (www.clsi.org) defines LoD as the lowest amount of a target that can be detected with probability but not quantified to an exact value. CLSI defines LoQ as the lowest amount that can be quantitatively determined with acceptable precision under controlled experimental conditions (Tholen et al., 2004; Forootan et al., 2017).

### 2.7. Protocol for Tier 3 (ddPCR) assay

Duplex ddPCR reactions (22 μL final volume) consisted of 11 μL ddPCR Supermix for Probes (2X) (No dUTP) (Bio-Rad Co., Berkeley, United States, Cat. #1863024), 7.9 μL of hpH_2_O, 2 μL of genomic DNA template, 1.1 μL of each of the primers at 900 nM concentration and 1.1 μL *Actin* and Y-chrom probes at 250 nM concentration. Variable number of NTCs were used as negative controls per assay session.

To avoid surpassing the upper limit of the dynamic range of the ddPCR reactions of 10^5^ copies of target DNA (Pinheiro et al., 2012; Podlesniy and Trullas, 2017), male *C. capitata* gDNA were diluted 10 times using a single 10-fold dilution from the original concentration for subsequent ddPCR reaction mixes since *Actin* and the Y-chromosome are both overexpressed in these male specimens and saturate the droplet generator and droplet reader above the dynamic range of the assay.

Twenty microliters of the duplex reactions were loaded in a QX200™ Droplet Generator to generate water-oil emulsion droplets (Bio-Rad Co., Hercules, United States) in eight-channel droplet generator cartridges per the manufacturer’s instructions. The resulting droplet mixes (∼40 μL) were then transferred into a 96-well PCR plate with low-retention pipette tips and sealed with PCR Plate Heat seal foil using a Bio-Rad PX1 PCR Plate sealer. Amplification in single water-oil emulsion droplets was performed *via* PCR in an Applied Biosystems^TM^ ProFlex^TM^ PCR System in 40 μL reactions with the following cycling conditions: 95 °C for 10 min followed by 40 cycles of 94 °C 30 s and 60 °C for 4 min, followed by one cycle of 98 °C for 10 min, and a ramp rate of 2.5 °C/s. Finally, the PCR plate was transferred into a QX200™ Droplet Reader and the results analyzed using Bio-Rad QX Manager Software 2.3.1 Standard Edition (Table 1 for summarized description of protocols).

Sensitivity limits for the ddPCR assay were determined using the following ddPCR assays:

For LoB, the highest number of false Y-chrom positive droplets potentially signaled with the assay, was calculated using 40 duplex ddPCR reaction replicates of a *v*♀ from Hawaii and a positive control ♂ from Hawaii (4 replicates), as well as a negative NTC (4 replicates). The two *C. capitata* DNA yields were ten-fold diluted to 10^-2^ with hpH_2_O. The LoB was calculated as LoB=Mean_blank_ + 1.645 × SD_blank_. Mean_blank_ corresponds to the mean number of false positive droplets detected in the 40 replicated reactions. We used a *v*♀ in order to detect the *Actin* control asserting the integrity of the specimen, while being negative for the Y-chrom. The assay considers a true positive if the number of copies exceeds the LoB (Supplementary Figure 1).

For LoD we used 40 duplex ddPCR reaction replicates of an SIT *C. capitata s*♂ after 10-fold serial dilutions of DNA yield to 10^-4^ with hpH_2_O. Positive control male *C. capitata* from Hawaii at 10^-3^ (4 replicates), as well as a negative NTC (4 replicates) were also used. LoD is calculated as LoD=LoB + 1.645 × SD_low concentration sample,_ as per CLSI Tholen et al. (2004) guidelines (Supplementary Figure 2).

For LoQ we used 40 ddPCR duplex reaction replicates of a *v*♀ DNA ten-fold diluted to 10^-1^, spiked with ten-fold dilutions of both *Actin* and Y-chrom synthetic amplicons ten-fold diluted to 10^-9^ using 1 μL of the DNA spiked mix per reaction. This mix ensured both specimen DNA and synthetic amplicon DNA could be tested to the lowest possible concentrations capable of being detected by the ddPCR technology, instead of all blank detections below these dilutions, which we tested. Positive control male *C. capitata* from Hawaii at 10^-3^ (4 replicates), as well as a negative NTC (4 replicates) were also used (Supplementary Figure 3).

The LoQ differs from LoD since detection is a qualitative measure of signal to noise detection ratio, whereas quantification is a quantitative measure of target detection, which is always higher than the qualitative measurement, as LoQ = LoD + (1.645 × SD_low concentration sample_).

## 3 Results

### 3.1 *C. capitata* gDNA extraction yields per trap type, locality and weathered time

Total gDNA retrieved from *C. capitata* specimens varied considerably depending on weathering time and conservation status of the specimens. Total gDNA yields (*Actin* + Y-chrom) in ML liquid traps decreased by a factor of 3 to 5 when compared to JK traps, with exception of Argentinian specimens at three weeks weathering in ML traps (Table 2). We retrieved more gDNA from females than males (Figure 1), most likely because the ovipositor adds an additional 1.2 mm to the abdominal length of females which also present a larger diameter to produce and store eggs (Weems, 1981; Plácido-Silva et al., 2005), hence more tissue from where to extract DNA. Differences in gDNA yields were significant between sexes and within or across treatments, with lower gDNA obtained from ML liquid traps and males (Figure 1).

**Table 2.** *C. capitata* total gDNA extraction yields (ng/µL) discriminated by country and sample type, for both sexes preserved in EtOH, JK traps, ML traps, and weathered times of zero weeks (Wks) to three Wks. Total gDNA quantified using a Qubit™ 4 Fluorometer. Mean and Standard deviation calculated using JMP® version 19.1, Statistical Discovery LLC, Cary, NC, USA.

| Country<br>Sample type | Argentina | Hawaii, USA | Guatemala |  |
| --- | --- | --- | --- | --- |
|  | Mean ± SD | Mean ± SD | Sample type | Mean ± SD |
| EtOH 0 Wks ♂ | 19.0 ± 0.8 | 17.3 ± 8.0 | EtOH 0 Wks s♂ | 8.1 ± 5.4 |
| EtOH 0 Wks m♀ | 20 ± 0.0 | 28.8 ± 1.25 |  |  |
| EtOH 0 Wks v♂ | 20 ± 0.0 | 18.4 ± 2.9 |  |  |
| JK 2 Wks ♂ | 22.7 ± 1.3 | 13.1 ± 5.2 | JK 2 Wks s♂ | 5.2 ± 2.8 |
| JK 2 Wks m♀ | 18.5 ± 2.5 | 13.4 ± 1.3 |  |  |
| JK 2 Wks v♂ | 21.8 ± 2.2 | 16.3 ± 5.8 |  |  |
| JK 3 Wks ♂ | 29.1 ± 1.8 | 8.5 ± 5.3 | JK 3 Wks s♂ | 31.7 ± 5.4 |
| JK 3 Wks m♀ | 25.7 ± 9.9 | 15.0 ± 6.6 |  |  |
| JK 3 Wks v♂ | 20 ± 0.0 | 23.8 ± 1.8 |  |  |
| ML 2 Wks ♂ | 4.8 ± 2.1 | 1.2 ± 0.3 | ML 2 Wks s♂ | 2.1 ± 2.0 |
| ML 2 Wks m♀ | 1.0 ± 0.2 | 8.2 ± 1.8 |  |  |
| ML 2 Wks v♂ | 4.96 ± 0.9 | 1.9 ± 1.3 |  |  |
| ML 3 Wks ♂ | 2.9 ± 1.3 | 1.5 ± 0.8 | ML 3 Wks s♂ | 4.1 ± 4.5 |
| ML 3 Wks m♀ | 20 ± 0.0 | 7.1 ± 5.6 |  |  |
| ML 3 Wks v♂ | 4.9 ± 3.7 | 21 ± 0.8 |  |  |

**Figure 1.**
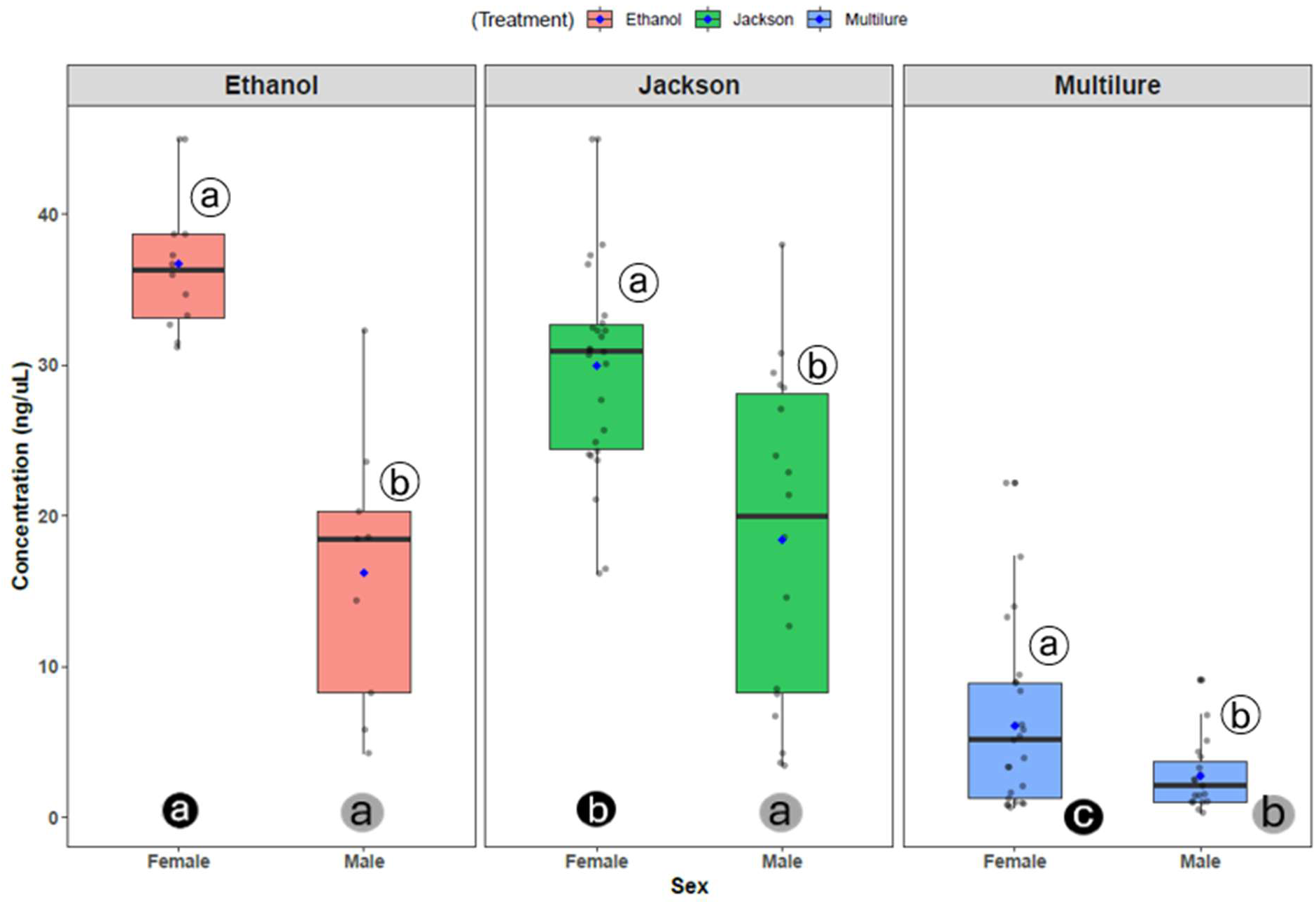
Total gDNA concentration (ng/µL) from *C. capitata* specimens by sex and treatment (Ethanol, JK and ML traps). Statistically significant differences between sexes within treatment type (white circled letters) and across treatment type (black circled letters comparing females and grey circled letters comparing males) using ANOVA, Tukey HSD post-hoc test, p < 0.05. Graphics and data analysis using R software (version 4.4.1; R-Core Team, 2026).

### 3.2 cPCR assay

The assay successfully detected the target bands, i.e. Y-chrom amplicon of 253 bp and *Actin* amplicon of 141 bp, across *C. capitata* specimen type, geographic regions, specimen condition and weathering time (Figure 2). Progressive decay of the JK dry traps specimens and ML liquid traps specimens in field conditions up to three weeks were sometimes detected, mostly in the ML specimens for the *Actin* amplicon. Background smears of lower molecular weight DNA could be observed and/or lower band intensity as well as non-specific amplification. However, this was

**Figure 2.**
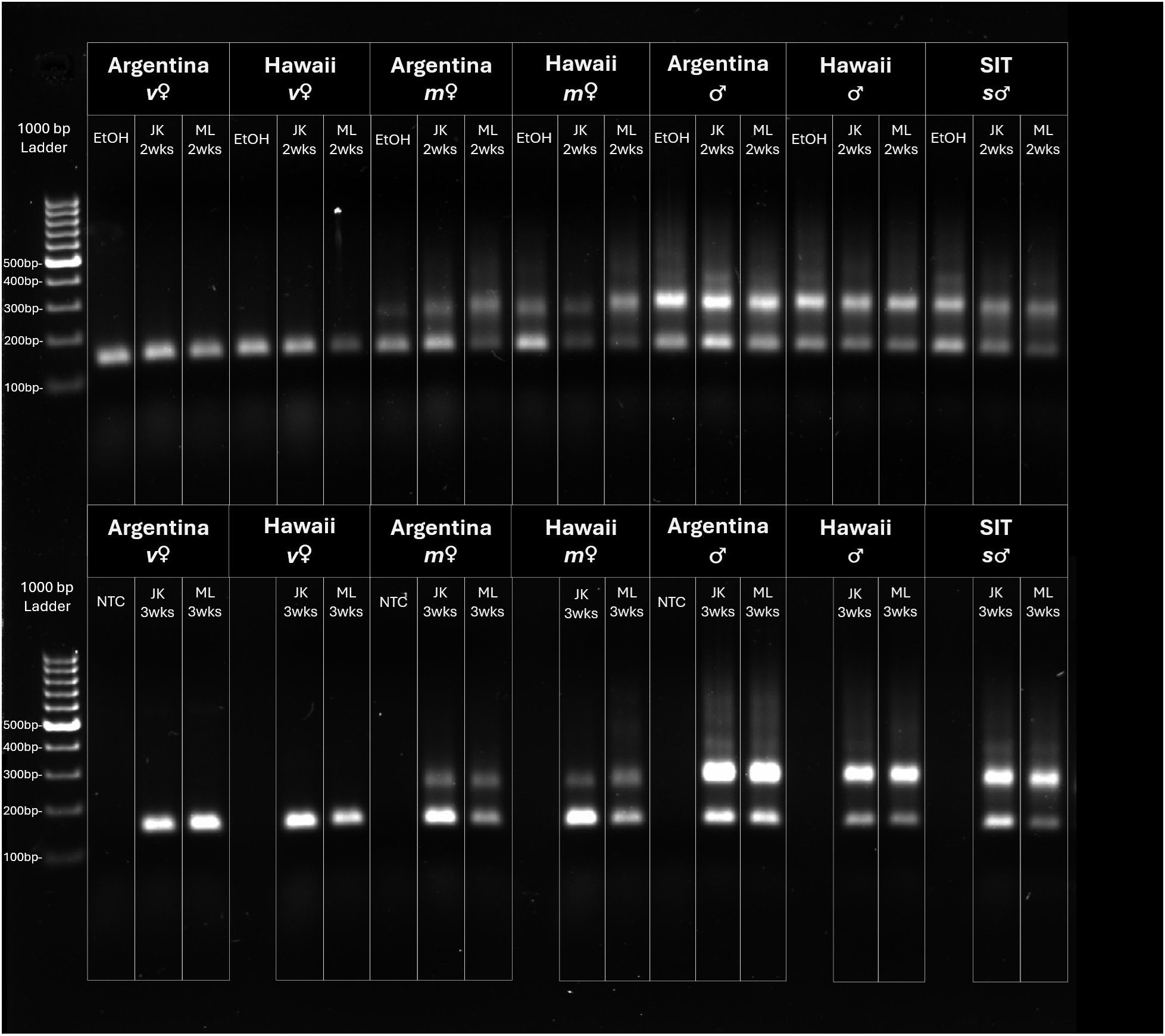
End-point conventional PCR gel electrophoresis visualization of target Y-chrom bands (253 bp bands) and *Actin* (141 bp bands) across geographic regions (Argentina, Hawaii, Guatemala SIT); specimen type (♂, *s*♂, *v*♀, and *m*♀) and weathering times in JK and ML traps. No Template controls (NTC) and 95% EtOH preserved specimens are also incorporated into the data.

marginal (∼10% of all samples) and did not compromise detection of target bands. Figure 2 depicts representative individuals for all sample types.

### 3.3 qPCR assay

#### 3.3.1 Analytical validation

The qPCR assay sensitivity and repeatability using synthetic amplicons of the target Y-chrom and *Actin* control genes in tenfold serial dilutions in both monoplex and duplex reactions displayed high linearity (R^2^ ≥ 99%) with amplification efficiencies ranging from 101.9% to 106.2%, thus validating a high qPCR performance with detection efficiencies as low as 11.7 copies/μL and 46.9 copies/μL for Y-chrom and *Actin*, respectively (Figure 3). In the duplex reaction 11.07 copies/μL for Y-chrom was detected in only one of three replicates, hence the reliable limit of detection for Y-chrom in the duplex reaction was 110.7 copies/μL with all three replicates being detected. *Actin* limit of detection in the duplex reaction was 46.9 copies/μL with all replicates being detected. For the monoplex reactions the detections were 11.7 copies/μL and 46.9 copies/μL for Y-chrom and *Actin*, respectively, and across all three replicates.

**Figure 3.**
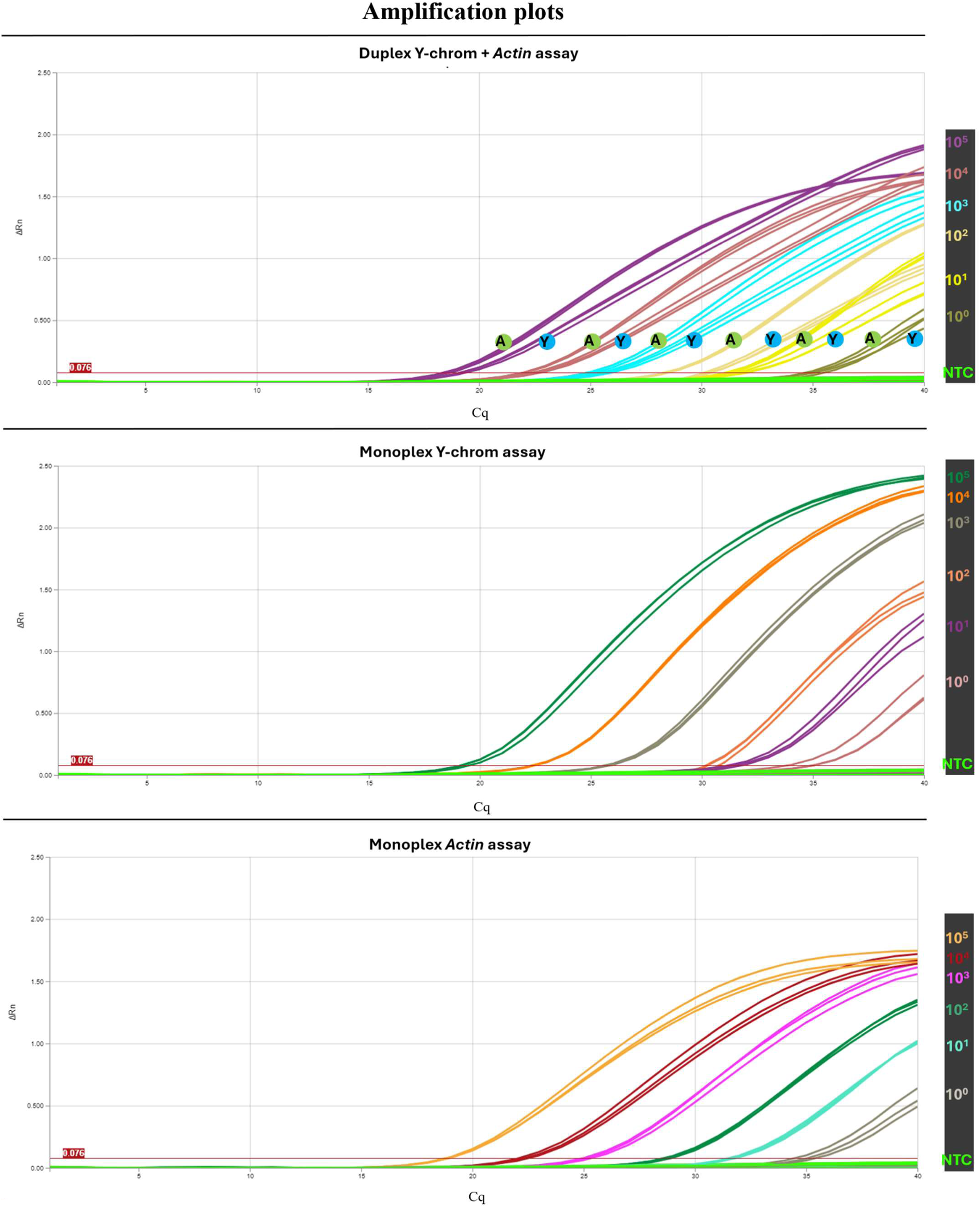
Duplex and monoplex amplification curves for the qPCR assay. Limit of detection of qPCR determined from serial ten-fold dilutions of synthetic amplicon template copies (from 1.107×10^5^ for Y-chrom (Y) and 4.695×10^5^ for *Actin* (A), to 11.7 and 46.9 copies/μL, respectively. Threshold for amplification detection (0.076) as per Thermo Fisher’s Design and Analysis 2 software default settings. Circled green A and Y blue identify the three replicates of the assay for a given template concentration of 10^5^ to 10^0^. All reactions were performed in triplicate. All assays run simultaneously in one qPCR assay run. Results were visualized using Thermo Fisher™ QuantStudio™ Design and Analysis 2 (DA2) software.

The assay displayed optimal PCR efficiency ranging from 101.9% to 103% in the duplex reactions for Y-chrom and *Actin* detection, respectively (Figure 4). Efficiency in the monoplex reactions was 106.2% to 104.6% for Y-chrom and *Actin*, respectively. Acceptable MIQE efficiency range is 90%-110% (Taylor et la., 2010) to account for random variability in pipetting or user precision (Schmidt et al., 2023; Bustin et al., 2025b).

**Figure 4.**
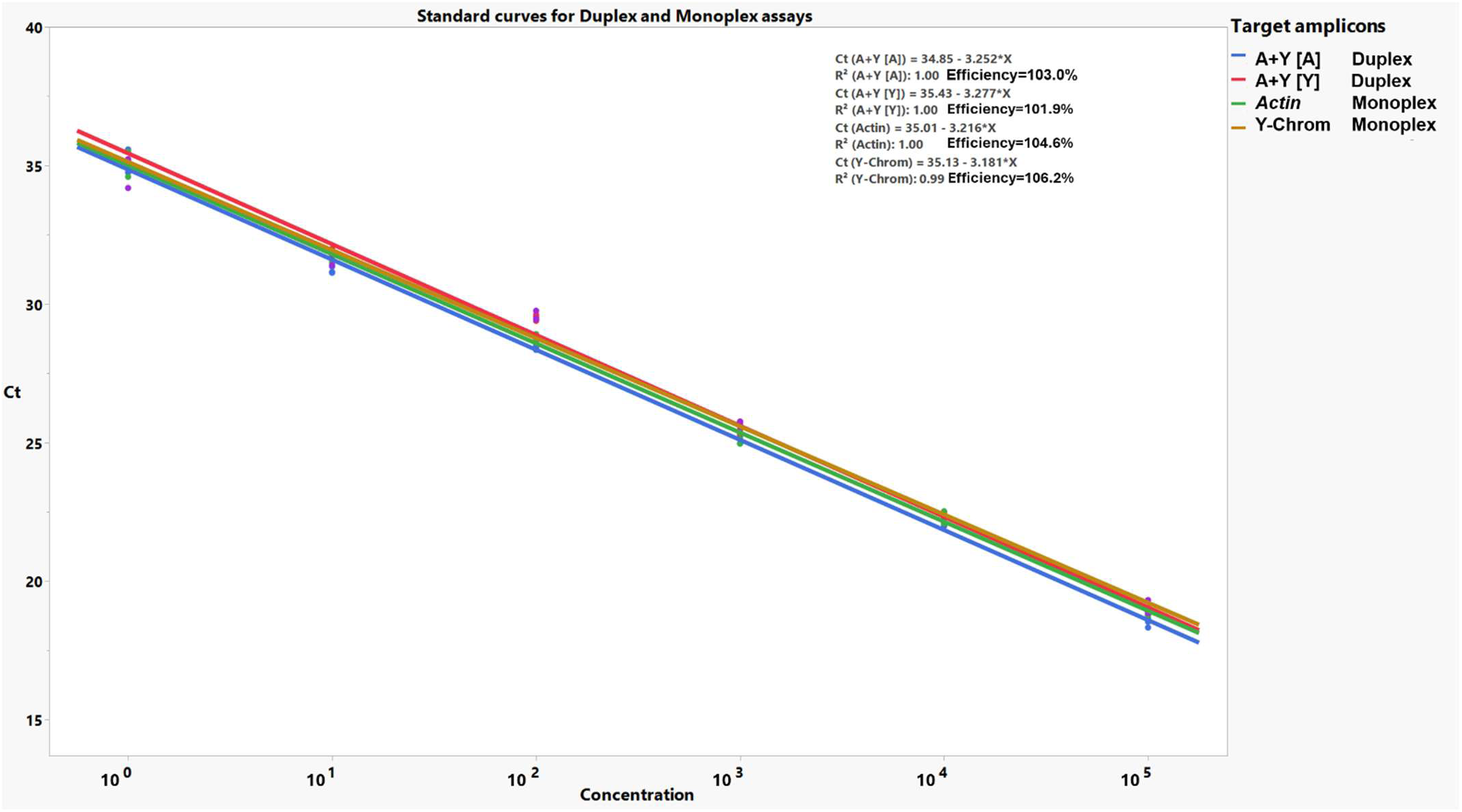
qPCR standard curve from the 10-fold dilution amplification curves in Figure 3. [A] and [Y] represent the standard curve for the duplex A+Y reactions, where A represents *Actin* and Y represents Y-chrom. Data depicts fitness to the regression line (R^2^). Efficiency = -1+10^(−1/slope)^ with the slope (average rate of change in probe detection for every decrease in amplicon concentration) in the regression equation as y=b_0_-b_1*_X, where b_1_=slope, as per MIQE guidelines (Bustin et al. 2025a). Efficiency was calculated using ThermoFisher Scientific qPCR Efficiency Calculator interface and data plotted using JMP® version 19.1, Statistical Discovery LLC, Cary, NC, USA).

#### 3.3.2 Sample validation

We tested ♂, *v*♀ and *m*♀ *C. capitata* colony specimens from Argentina and Hawaii, as well as SIT *s*♂ from Guatemala, in ML and JK traps, weathered for 2-and-3 weeks in the traps. In addition, we also tested fresh specimens preserved in EtOH 95% as positive controls. Efficiency of the qPCR was 100% across samples (See Supplementary Table 1 and Figure 5 A-C) for either target Y-chrom amplicon and *Actin* housekeeping control gene, thus attesting for both the presence of the Y-chrom in *m*♀, as well as the physical integrity of the specimens with the *Actin* housekeeping gene amplicon. For some of the samples the detection was at the accepted limit of detection threshold of Cq=35 cycles for Y-chrom or *Actin* (Supplementary Table 1). This mostly occurred in samples exposed for 2 or 3 weeks in 10% PG ML traps. These traps were particularly detrimental for the preservation of the physical integrity of the specimens, with *Actin* being mostly detected at *circa* 5 cycles after Y-chrom detection and with a lower fluorescence signal compared to dry traps (Figure 5A). This effect was less pronounced in *m*♀, although the fluorescence signal remained lower than the Y-chrom amplicon (Figure 5B).

**Figure 5.**
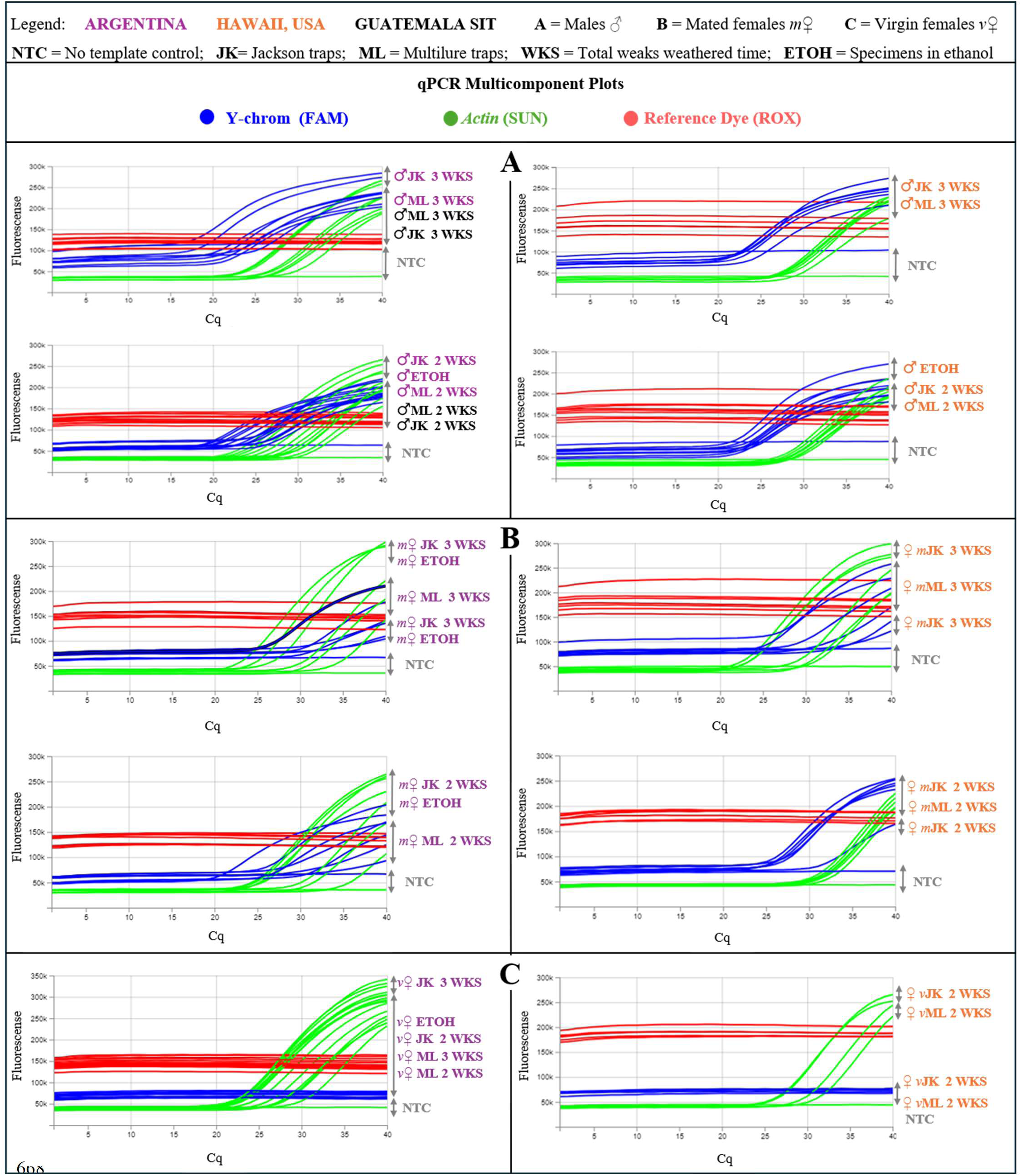
Validation of qPCR across geographic regions (Argentina, Hawaii, Guatemala SIT); specimen types (♂, *v*♀, *m*♀, Guatemala *s*♂), and weathering times, in JK and ML traps. NTCs and EtOH preserved specimens were also incorporated into the assays and data analysis. Amplification profiles display the Cq detection cycle and fluorescence signal amplitude for any given sample type and target (Y-chrom) or housekeeping gene (*Actin*), while flat lines represent no detection of target DNA amplicons. Sex and mating status results grouped per panel as males (**Panel A**), mated females (**Panel B**), virgin females (**Panel C**). Results were visualized using Thermo Fisher™ QuantStudio™ Design and Analysis 2 (DA2) software.

Total gDNA concentration (ng/μL) in ML liquid traps corroborates the lower amplification signals, and later Cq cycle detection in qPCR observed for *Actin* in these traps, with a significant lower yield of gDNA compared to EtOH or JK traps. Dry JK did not significantly reduce the total gDNA yields when compared with the EtOH preserved specimens (Figure 1).

All samples were subsequently re-tested and reconfirmed for mating status with the ddPCR Tier 3 assay.

### 3.4 ddPCR assay (Tier 3)

#### 3.4.1 Analytical validation

The detection resolution power and precision of our ddPCR assay yielded an LoB=0.43 copies of the Y-chrom target amplicon, in a 22 μL reaction; an LoD=4.44 copies per reaction, and a LoQ=14.31 copies per reaction with 95% probability (See Supplementary Fig 1-3). This precision was obtained from 40 replicates reactions per assay, as listed in the methodology (Supplementary Figure 1-3). We determined the specific dynamic range of our ddPCR assay as 10^3^ copies per μL (Figure 6), which is defined as the maximum amount of input target amplicons copies per microliter that can be quantified without loss of correlation between input and ddPCR detection. Above 10^3^ copies per μL correlation is not optimal (10^5^ copies per μL ddPCR assay can be consulted in the GitHub online repository for this manuscript).

**Figure 6.**
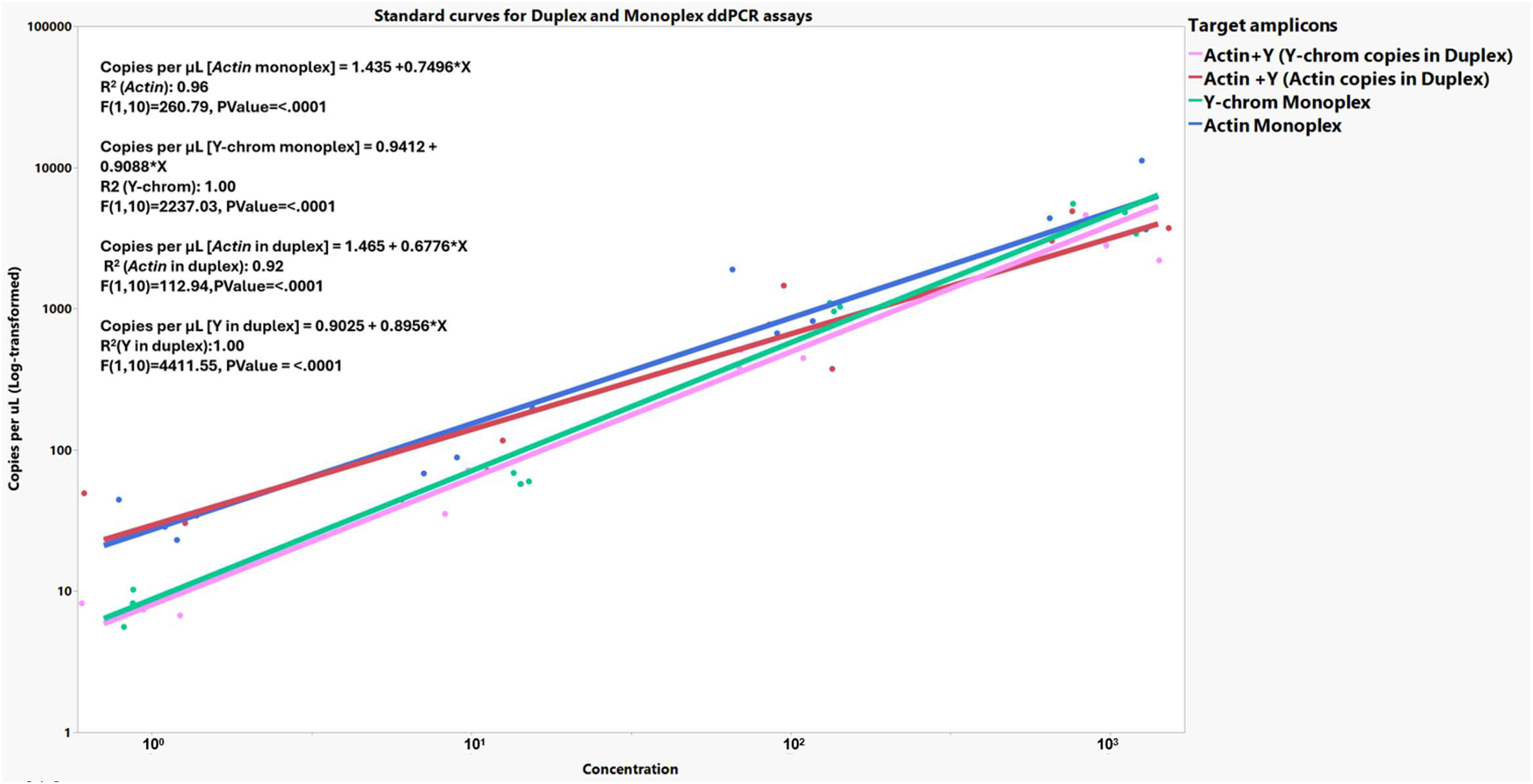
Dynamic range of the ddPCR assays quantifying the correlation between number of copies per μL in ddPCR monoplex and duplex reactions for both target amplicons. Y-chrom and *Actin*, against 10-fold serial dilution of synthetic target amplicons based on three replicates per dilution. Regression equation as y=b_0_-b_1*_X, where b_1_=slope. The fitness to the regression line determined as R^2^, and the correlation of copies of target amplicons as concentration rises is determined as significant at a *p*-value of *p* < 0.0001. Graphics using JMP software v. 19.1, JMP® Statistical Discovery LLC, Cary, NC, USA.

#### 3.4.2 Sample validation

Absolute specificity of the ddPCR for the Y-chrom probe detection was confirmed across all sample types and weathering times (Supplementary Table 1, Figure 7A-C) with 100% for *m*♀, ♂ and *s*♂ testing positive for Y-chrom and *Actin* control genes, independently of their preservation status and geographic origin, whereas *v*♀ tested negative for Y-chrom, and positive for *Actin*.

**Figure 7.**
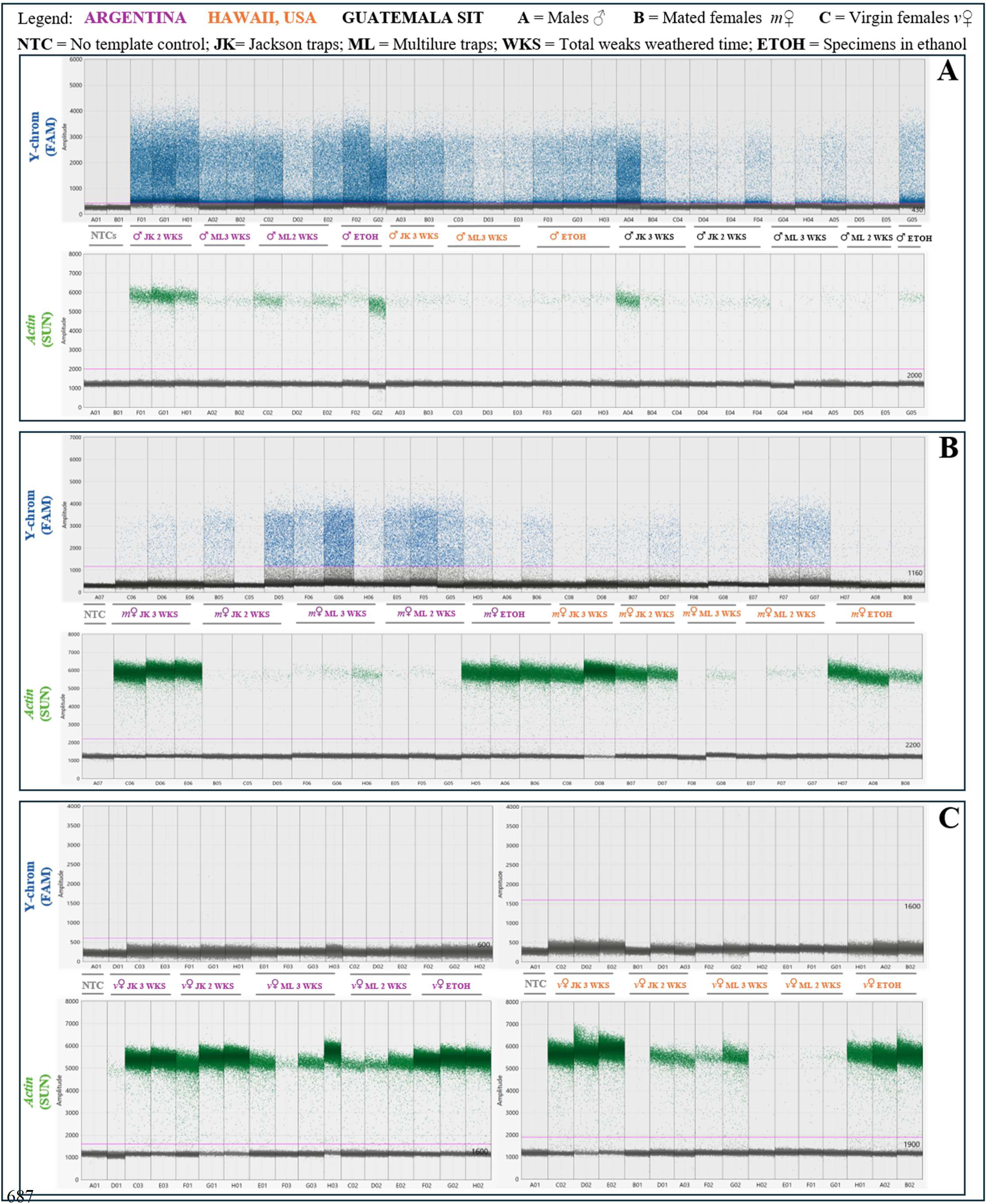
Validation of ddPCR across geographic regions (Argentina, Hawaii, Guatemala SIT); specimen type (males, virgin females, mated females), and weathering times in JK and ML traps. Sex and mating status results grouped per panel as males (**Panel A**), mated females (**Panel B**), virgin females (**Panel C**). NTCs and 95% EtOH preserved specimens were also incorporated into the dataset. Digital droplet profiles display the amplitude of each droplet generated for a given sample. Samples are coded per droplet column labelling both the above (Y-chrom) and below chart (*Actin*) for each plot and panel. The droplet with the highest amplitude observed in each NTC control was used as the threshold (in pink) for positive vs. negative detection of the Y-chrom and *Actin* targets. Black droplets represent negative droplets lacking the target DNA amplicons while colored ones represent target detection. Green for *Actin* and blue for Y-chrom. Results analyzed with Bio-Rad QX Manager Software 2.3.1 Standard Edition

Positive droplets and fluorescence amplitude varied considerably depending on the conservation status of the sampled individuals, with specimens retrieved from ML liquid traps displaying fewer positive droplets for *Actin*, hence lower physical integrity (Supplementary Table 1, Figure 7A-B). This finding concurs with the same observation in the qPCR results. Y-chrom positive droplets also varied, although less impacted by the decline in physical integrity of specimens in our data, but more plausibly as a reflection of the variable amount of sperm present if the *m*♀ given that ♂ and *s*♂ courtship behavior in *C. capitata* often involves intermittent, successful or not, mount and copula attempts which directly affect transferred yields of sperm (Prokopy and Hendrichs, 1979). This behavior is particularly notorious in lab reared specimens, such as ours, with a more limited capacity to copulate with females and more intermittent attempts to do so (Briceño et al. 2007).

Positive droplets ranged from saturation (notably in ♂ and *s*♂,) to scarce droplets, although in the LoQ range of the assay (Supplementary Table 1, Figure 7A). These observed droplets were all within the dynamic range of our ddPCR at 10^3^ copies per reactions. For the Y-chrom detection we observed limited droplet separation between positive and negative droplets, however still distinguishable by Bio-Rad’s QX Manager software using the NTC samples as the threshold baseline. Manual thresholds are more reliable and preferred for complex assays with low abundance of target amplicons in order to ensure absolute quantification accuracy. In addition, duplex ddPCR enhances assay complexity with the high-abundance amplicon being preferentially sequenced, and both primer sets and probes sharing the same DNA polymerase and chemistry in reaction (Elnifro et al., 2000). We observed this complexity at play with Y-chrom vs. *Actin* being detected preferentially depending on the preservation status of the specimen in ML liquid traps with *Actin* preservation diminished, hence, *m*♀ Y-chrom detection enhanced (Figure 7B), whereas dry traps (JK) with better *Actin* preservation, as well as EtOH preserved samples, displayed stronger *Actin* detection and lower Y-chrom detection (Figure 7B). This observation is also confirmed in most of the ♂ and *s*♂ (Figure 7A).

Although we explored different concentrations of primers and probes, as well as different annealing temperatures, we determined that the default concentrations suggested by the QX200 manufacturer at 250/900 nM probes/primers and annealing temperatures gradients below or above 60 °C did not provide better separation. Extension time increase in the PCR step also did not increase separation. Amplicon lengths exceeding 200 bp pose challenges for amplification. This coupled with the presence of tandem repeat AT enriched regions up to 83% in the Y-chrom of *C. capitata* (Zhou et al., 2000), as the Y-chrom specific amplicon we used, could account for the observed reduction in the amplitude and separation between negative vs. positive droplets. Nevertheless, no *v*♀ or *m*♀ were incorrectly assigned using our LoQ parameters.

We also determined the concentration of target Y-chrom and *Actin* amplicons detected by our ddPCR duplex assay to further determine the impact of JK dry and ML liquid traps, as well as sex and ♂ and *s*♂ status, on DNA detection (Figure 8A-C). This can serve as a measure of the precision of our ddPCR assay to detect the targets genes, even with decayed specimens, or irradiated specimens, over the course of a maximum of three weeks weathering time and unpredictable environmental conditions. We determined that the average *Actin* detected (Figure 8A) did not vary significantly across trap, sex and specimens preserved in EtOH. However, the traps displayed a broader range of concentrations while the specimens in EtOH were more consistent. ML traps affected the amount of *Actin* detected for females, although not statistically significant when compared with JK (Figure 8A). For the Y-chrom, the variability of concentrations detected in ♂ and *s*♂ was broader in the traps (Figure 8B) and weathered weeks (Figure 8C) than the one observed for Actin (Figure 8A). Notably *s*♂ Y-chrom detection was much lower than the fertile ♂. SIT genetic sexing strains use controlled radiation to render specimens infertile while causing translocation mutations in the Y-chromosome (Porras et al., 2020), therefore the lower detection is plausibly correlated with the impaired genetic nature of these specimens. Week appeared to further impact Y-chrom detection as weathering time increased from 2 to 3 weeks, with *s*♂ yielding less gDNA than fertile ♂ (Figure 8C). In fact, for three-week weathering times the detection of the Y-chrom is superior to week two. Three-week field weathered samples undergo more biological breakdown than two-week weathered samples. This decay might act as an enzyme cleaving the bounded complementary strands in the double-stranded DNA (dsDNA) material of the specimen. During ddPCR droplet generation the pre-cleaved single DNA strands (ssDNA) will be partitioned and encapsulated into two different droplets, which in turn will generate two signals during PCR, rather than one signal in well preserved dsDNA encapsulated in single droplets. This biological artifact is documented and does not impact accuracy of detection (Kline et al., 2020; Kim et al 2026), but inflates the detection parameters. The sample decay is inevitable in our source material, however our assays could successfully overcome these challenges with accurate detections of target Y-and-*Actin* amplicons.

**Figure 8.**
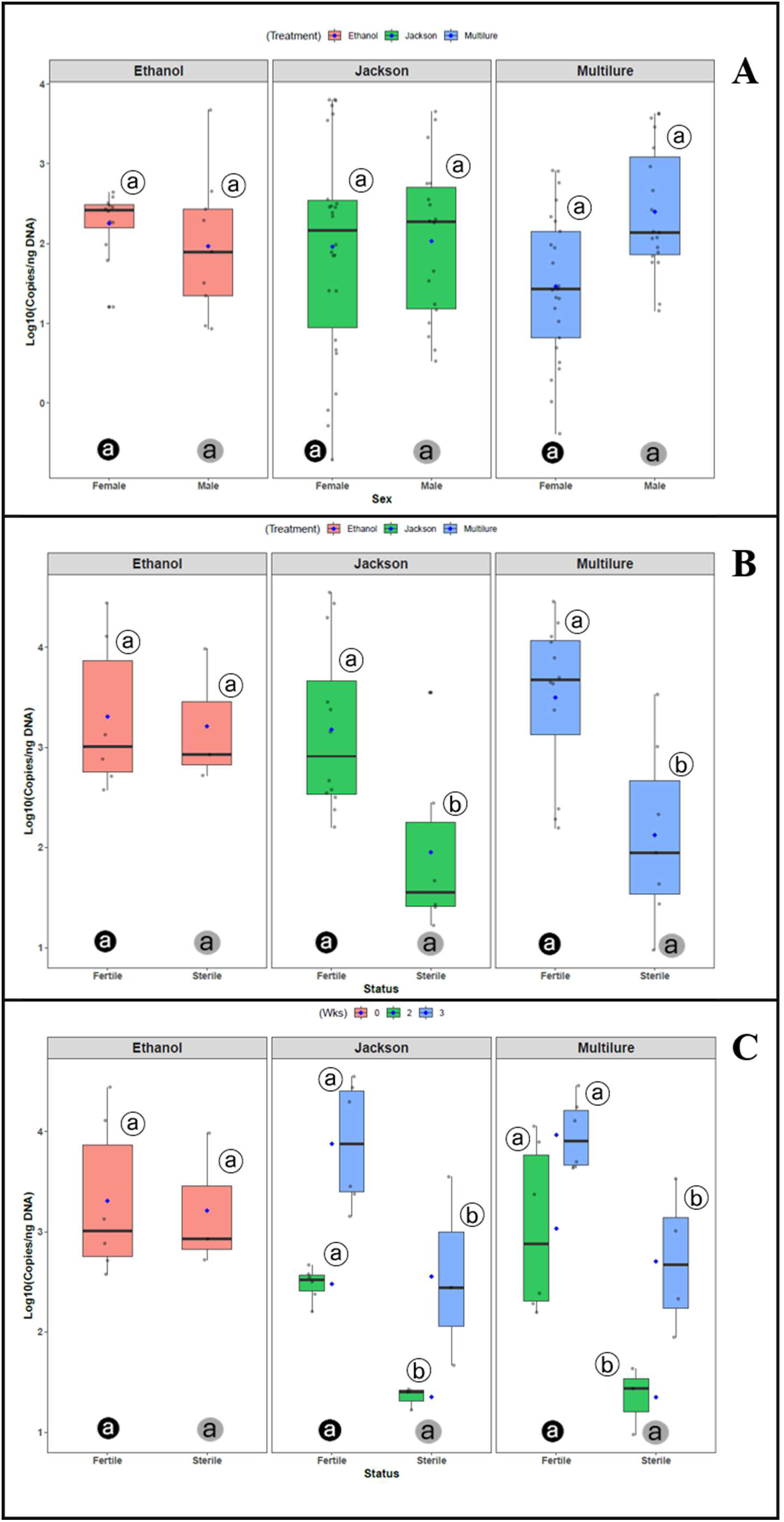
Depiction of the concentration of target Y-chrom and *Actin* amplicons as number of copies per µL detected by our ddPCR duplex assay as a function of treatment (EtOH, JK and ML traps), sex and ♂ and *s*♂ irradiation status. **Panel A** displays *Actin* detection per sex, EtOH and trap type. **Panel B** displays Y-chrom detection as a function of irradiation status, as fertile ♂ or irradiated sterile males, *s*♂. **Panel C** displays Y-chrom detection per weathering time, i.e., two weeks (Wks) or three Wks, as well as EtOH and trap type for ♂ vs. *s*♂. Statistically significant differences between sexes or irradiated status, within treatment type (circled white letters) and across treatment types (dark circled letters comparing females and grey circled letters comparing males for panels A and B, and fertile ♂ vs. *s*♂ for Panel C) using ANOVA, Tukey HSD post-hoc test, *p* < 0.05. Graphics and data analysis using R software (version 4.4.1; R-Core Team, 2026).

## 4 Discussion

The three-Tier DNA detection method demonstrated excellent analytical performance including precision, linearity, and detection capability, increasing with Tier level from conventional end-point PCR to droplet digital ddPCR. This approach overcame potential issues from limited preservation of field specimens retrieved in JK dry traps and ML liquid traps after 2- and 3-weeks, hence validating the practical use of the assay detecting mating status of females in monitoring/surveillance grids for Medfly in the United States of America, and beyond.

The increasing resolution power of the multi-tiered Y-chrom detection method proposed here allowed first, a quick preliminary screening of sample physical and genetic preservation status and mating status *via* cPCR; second, a quantification and real-time detection of sperm presence *via* qPCR in *m*♀; and third, detection of minimal traces of sperm in *m*♀ *via* ddPCR. Sensitivity reached 100% for both qPCR and ddPCR. Performance below 100% accuracy was only observed at Tier 1, *via* standard end-point PCR and gel electrophoresis which lacked *Actin* detection in two specimens (see Supplementary Table 1A). Preservation in traps, especially in ML liquid traps such as these two specimens, can hinder the performance of conventional PCR and, as we have shown, impacts the physical integrity and the amount of useable DNA for downstream applications. The samples listed above were then processed with qPCR and ddPCR which enhanced accuracy and precision detecting the Y-chrom and *Actin* amplicons (Supplementary Table 1A), demonstrating that this methodology can overcome the quick physical degradation of the minute *C. capitata* in JK dry and ML liquid field traps.

Conventional PCR poses known technical limitation in reproducibility given the required in-hand laboratory manipulation in pre-and-post amplification steps and does not display tolerance to PCR inhibitors in reaction (Yang and Rothman, 2004). This coupled with the unavoidable degradation of the Medfly samples in the traps, and given cPCR is a qualitative measurement of DNA, rather than quantitative as qPCR and ddPCR, reliability on the integrity of specimens is more pressing for cPCR. Moreover, cPCR requires significant more genomic DNA than qPCR and ddPCR (Engleberg, 1992), which is our case can be scarce for the target Y-chrom amplicon in *m*♀. Finally, the multiplexing of the cPCR in our assays with target Y-chrom primers and *Actin* control gene primers also posed known challenges for cPCR (Polz et al., 1997; Elnifro et al, 2000), mostly in the preferential amplification of *Actin* or Y-chrom depending on the integrity of the specimen. We opted to include cPCR as an initial preliminary screening of sample quality and target amplicon presence given the feasibility of most molecular laboratories to perform cPCR and given that our assay performance is still exceptionally high for cPCR, at 98% accuracy for *Actin* and 100% accuracy for Y-chrom across samples. The assays in the more sensitive platforms qPCR and ddPCR, overcame cPCR limitations with an accuracy and precision of 100% detection of the target amplicon in ♂, *s*♂, *v*♀ and *m*♀ independently of the conservation and tissue integrity of the specimens, in weathered condition up to three weeks.

Based on our results, we strongly recommend for regulatory purposes to test samples via qPCR. If detection of Y-chrom in *m*♀ occurs at Cq ≥35, ddPCR should be performed to confirm the presence of the Y-chrom at low concentrations. We also strongly recommend the proper conditioning of specimens during transit to laboratory facilities and alternative preservation liquids for ML liquid traps given the observed decline in the physical integrity of the specimens for the *Actin* muscle gene. Commercial formulation of eco-safe PG antifreezes, commonly used in these traps, also incorporate other chemicals apart from PG, namely colored dyes, which might interfere with the physical preservation of the specimens.

The three-tier method presented in this manuscript offers an unambiguous and unequivocal confirmation of mating status of female *C. capitata*, in opposition to DAPI staining techniques with noticeable limitation such as technical and taxonomic skills required to dissected specimens and locate the target tissues and cells for staining, apart from the mutagenic toxic nature of the product. The three-tier methodology proved successful with increasing detection and quantification power of the Y-chrom in *m*♀, with a LoQ of 110.7 copies/μL in qPCR and 14.31 copies/reaction in ddPCR, and a LoQ of 4.4 copies/reaction in ddPCR which translates to just 14 spermatozoa in a *m*♀. The methodology can determine mated status of *C. capitata* eliminating any subjectivity when determining the need to initiate extensive quarantines and multi-million eradication programs.

## 5 Data Availability Statement

The bioinformatic data files used to validate qPCR and ddPCR assays for this study can be found at GitHub online repository under https://github.com/Jmar-06/3_Tier_assay-Medfly-mating-status.git. Any further queries should be directed to the corresponding author.

## Authors contributions

JAPM and CJZ designed and executed the study. JAPM, HU, MRM contributed to investigation, data acquisition and formal analyses. MSS and AH implemented mating arenas and provided *C. capitata* specimens of both sexes and mating status for the study. KF and JS supervised study development and project administration. JAPM and CJZ wrote the first draft, subsequently edited and approved by all co-authors.

## Funding

This research was funded by the Florida Department of Agriculture and Consumer Services, Division of Plant Industry.

Mention of trade names or commercial products in this publication is solely for the purpose of providing specific information and does not imply recommendation or endorsement by FDACS-DPI and the USDA. FDACS-DPI and the USDA are equal opportunity employers.

## Acknowledgements

We thank the Florida Department of Agriculture and Consumer Services, Division of Plant Industry (FDACS-DPI), for their support of this work, as well as FDACS-DPI staff Lynn Combee, Cheryl Roberts, Hannah Luciano, Paola Dixon, Craig Welch and Matthew Meise. Also, Mark Spearman and Jason Spiller at the United States Department of Agriculture Sterile Insect Release Facility (USDA-SIRF) for providing euthanized *C. capitata* specimens of SIT irradiated male Medfly. We also thank Alejandro Asfennato and Diego de Rosas, respectively at Instituto de Sanidad y Calidad Agropecuaria Mendoza (ISCAMEN), and Susbin S.A., Mendoza, for providing specimens of *C. capitata* from Argentina.

