## Supplementary Figures 1 to 3 for "PCR–based assays for determining mating status in field–weathered *Ceratitis capitata* with enhanced precision across conventional, quantitative, and droplet digital platforms"

Supplementary Figure 1 – ddPCR Limit of Blank (LoB) calculated as LoB=Mean_blank_ + 1.645 × SD_blank_ as per CLSI Tholen et al. (2004) guidelines based on 40 duplex ddPCR reaction replicates plus positive (+) and negative (−) controls. Mean_blank_ corresponds to the mean number of false positive droplets detected in the 40 replicated reactions.

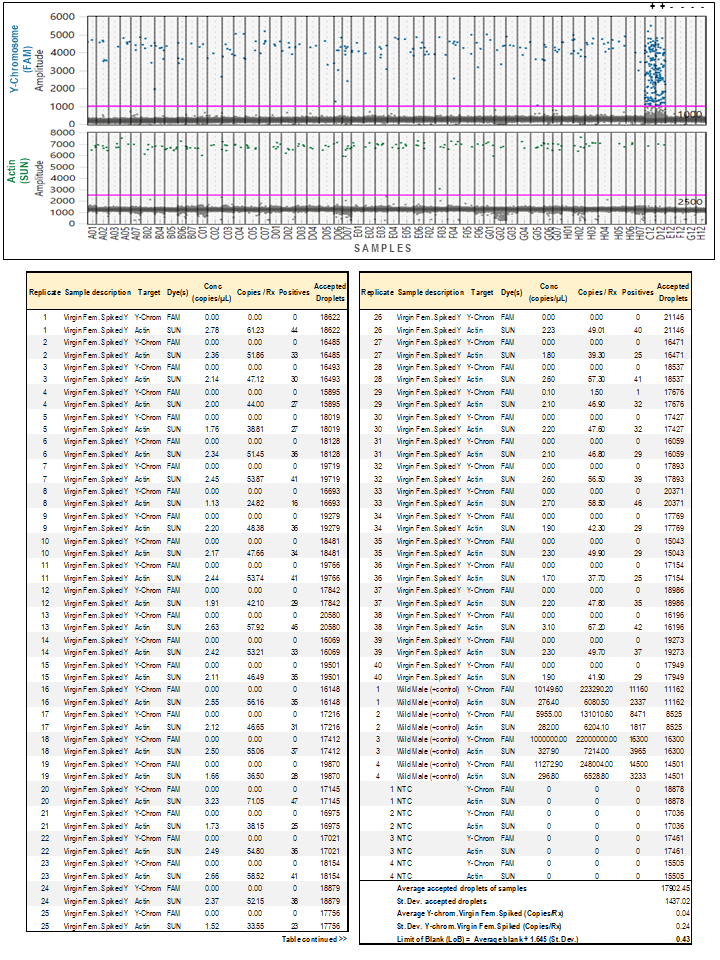

Supplementary Figure 2 – ddPCR Limit of Detection (LoD) calculated as LoB=LoB + 1.645 × SD_Low concentration sample_ as per CLSI Tholen et al. (2004) guidelines based on 40 duplex ddPCR reaction replicates plus positive (+) and negative (−) controls.

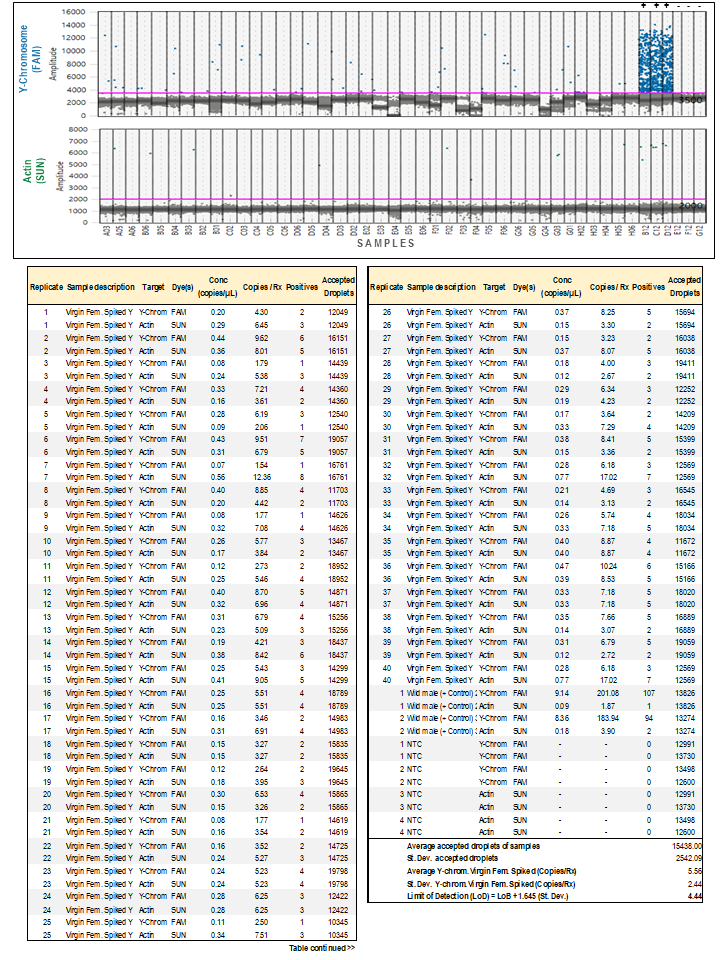

Supplementary Figure 3 – ddPCR Limit of Quantification (LoQ) calculated as LoQ=LoD + 1.645 × SD as per CLSI Tholen et al. (2004) guidelines based on 40 duplex ddPCR reaction replicates plus positive (+) and negative (−) controls.

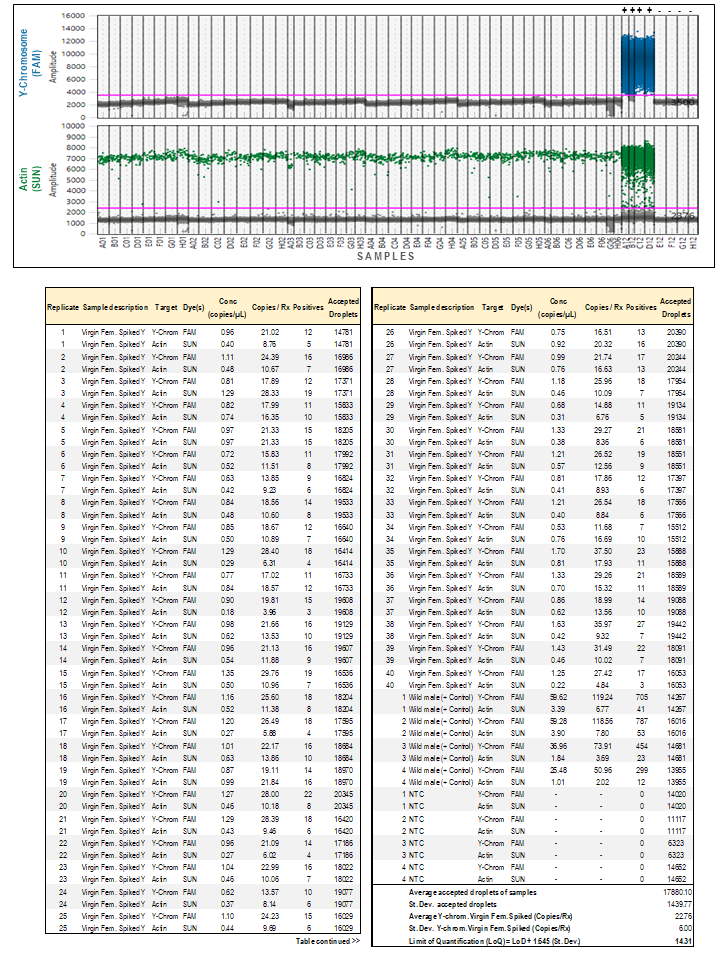
